# Transgenerational effects of exercise on mouse brain and cognition

**DOI:** 10.1101/2023.03.08.531840

**Authors:** Elisa Cintado, Patricia Tezanos, Manuela De las Casas, Pablo Muela, Kerry R. McGreevy, Ángela Fontán-Lozano, Eva Sacristán-Horcajada, Jaime Pignatelli, María L. de Ceballos, María Jesús del Hierro, Julia Fernández-Punzano, Lluis Montoliu, José Luis Trejo

## Abstract

Lifestyle induces long lasting effects on brain and cognition, with some interventions like stress including transgenerational inheritance mediated by epigenetic mechanisms. Physical exercise is one lifestyle intervention driving robust improvements of cognition, including intergenerational transmission to the litter. However, little is known about whether exercise effects are transgenerationally transmitted. Here we analyzed adult hippocampal neurogenesis (AHN) and behavioral phenotype of sedentary adult male mice of F2 generation of exercised grandfathers (F0). Both F1 and F2 were sedentary, while F0 performed moderate exercise. We found F2 mice from exercised F0 acquired and recalled both spatial and non-spatial information better than F2 from sedentary F0. Contextual fear conditioning was not affected, together with no differences in AHN markers. Hippocampal smallRNAseq analysis revealed 35 significant differentially expressed (sDE) microRNAs (miRNAs) associated to relevant brain function families. Moreover, 11 of the 35 miRNAs target gene sets were found also enriched in F0 and F1, as well as target genes of 6 of them were differentially expressed also in F0 or F1. One of these 6 is miRNA-144, that together with miRNA-298 were found inversely correlated to cognitive index in F2. These results demonstrate that transgenerational transmission of the effects of exercise on specific cognitive tasks persists after two generations, even though some cellular changes induced in F1 vanish in F2. Thus, they suggest moderate exercise training has longer-lasting effects than previously thought, probably mediated by a small group of miRNAs acting across generations, and this is worth taking into account in public health programs.

## INTRODUCTION

The beneficial effects of exercise on body and brain health are very well known (Trejo et al., 2002)(Llorens-Martín et al., 2008)(Di Liegro et al., 2019)(Umegaki et al., 2021)(Hashimoto et al., 2021)(McGillivray et al., 2021)(Meli et al., 2021)(Sutkowy et al., 2021)(Cantelon & Giles, 2021)(Mellow et al., 2022), as well as the adverse effects of a sedentary lifestyle (Martin et al., 2018)(Song et al., 2018)(Rendeiro & Rhodes, 2018). In fact, lifestyle interventions are known as potent drivers of neural plasticity in all type of organisms (Mora, 2013)(Valero et al., 2016)(Phillips, 2017)(Maharjan et al., 2020). Besides, the heritability of these lifestyle effects to either the next generation (intergenerational inheritance) or the second and later generations (transgenerational inheritance (Jirtle & Skinner, 2007)) has been largely explored in different taxa (Baugh & Day, 2020)(Frolows & Ashe, 2021)(Grishok, 2021)(Chey & Jose, 2022)(Zare et al., 2018)(Deas et al., 2019)(Hoyer-Fender, 2020)(Mu et al., 2021). It has also been explored in different models, with a special focus in stress or diet (Blaze & Roth, 2015)(Hime et al., 2021)(Lagisz et al., 2014)(Risal et al., 2021). However, the inheritance of the exercise-induced effects has been only recently addressed and mostly in patrilineal, intergenerational designs (Short et al., 2017)(Benito et al., 2018) (McGreevy et al., 2019)(Yeshurun & Hannan, 2019)(Yang et al., 2021).

The inter- and transgenerational effects of stress have been largely described (Gapp et al., 2014)(Jawaid et al., 2018)(Yuan et al., 2016)(Ambeskovic et al., 2020)(Ambeskovic et al., 2019). The evidence of direct and indirect consequences of stress and the mechanisms mediating its inheritance on next generations have paved the way to analyze the potential positive effects of other lifestyle interventions, as well as to set a gold standard of the methodology to address these questions in mice models (Bohacek & Mansuy, 2015).

As for physical exercise, most studies on intergenerational inheritance have focused on maternal exercise during pregnancy (Kusuyama et al., 2020)(Eclarinal et al., 2016). This germline-mediated matrilineal epigenetic inheritance is not easily distinguished from a non-germline epigenetic inheritance, mainly because the role of maternal intervention cannot be ruled out (Champagne, 2008). On the other side, we now know that positive, pro-cognitive effects, and the pro-neurogenic effects of moderate physical exercise in male mice are inherited by the males of the first generation (McGreevy et al., 2019)(Vieira de Sousa Neto et al., 2021). Specifically, it has been reported that animals of these litters show better spatial and non-spatial cognitive abilities, increased adult hippocampal neurogenesis, along with changes in brain mitochondria activation and hippocampal gene expression, including differential expression of target genes of several microRNAs like for example the 212/132 cluster in the hippocampus of both fathers and litters (McGreevy et al., 2019). Similar pro-cognitive results were reported after paternal environmental enrichment inherited by the first generation (Benito et al., 2018). In the latter study, the involvement of the microRNA 212/132 cluster, present in the male sperm of enriched animals, was elegantly demonstrated to be one mediating mechanism of the intergenerational transmission (Benito et al., *op.cit.*).

In this work we address whether the exercise-induced effects in the parental (F0) and first-generation litter (F1) are also transmitted transgenerationally to the second generation (F2). This is a relevant question both because, to our knowledge, there is no evidence for the transgenerational transmission of positive outcomes of lifestyle interventions focused on brain and cognition, and because this information would allow to design more impactful public health policies for the population affected by the sedentary life pandemic (Guthold et al., 2018).

## RESULTS

We analyzed here through a patrilineal design the behavioral phenotype and the AHN of sedentary F2 adult male mice, bred from sedentary F1 animals by *in vitro* fertilization (IVF) and embryo transfer (ET), which in turn were bred either from sedentary or exercised F0 mice by IVF and ET (Figure 1). We found a similar, normal exploratory activity in both groups as measured by an automated actimeter, showing no significant differences neither in horizontal (Figure 2a) nor in vertical (Figure 2b) activity, no differences in total distance covered (Figure 2c) or time in movement (Figure 2d). No differences were found in the time spent exploring the margin or the center of the cage (Figure 2e). The time course displayed an expected decrease both along the time during day 1, and a decrease in day 2 compared to day 1 in both groups (F2SED paired sample t-test t(14)=9.588 p<0.01; F2RUN paired sample t-test t(7)=5.856 p<0.01). It is noteworthy that the exercised F2 showed a significant lower activity than sedentary F2 animals during specific time intervals on day 2, when activity is segmented by minutes (Figure 2f, *post-hoc* comparisons corrected by Bonferroni minute 2 p=0.026, minute 4 p=0.049). This suggests F2RUN mice recognize the arena as a familiar context.

**Figure 1.**
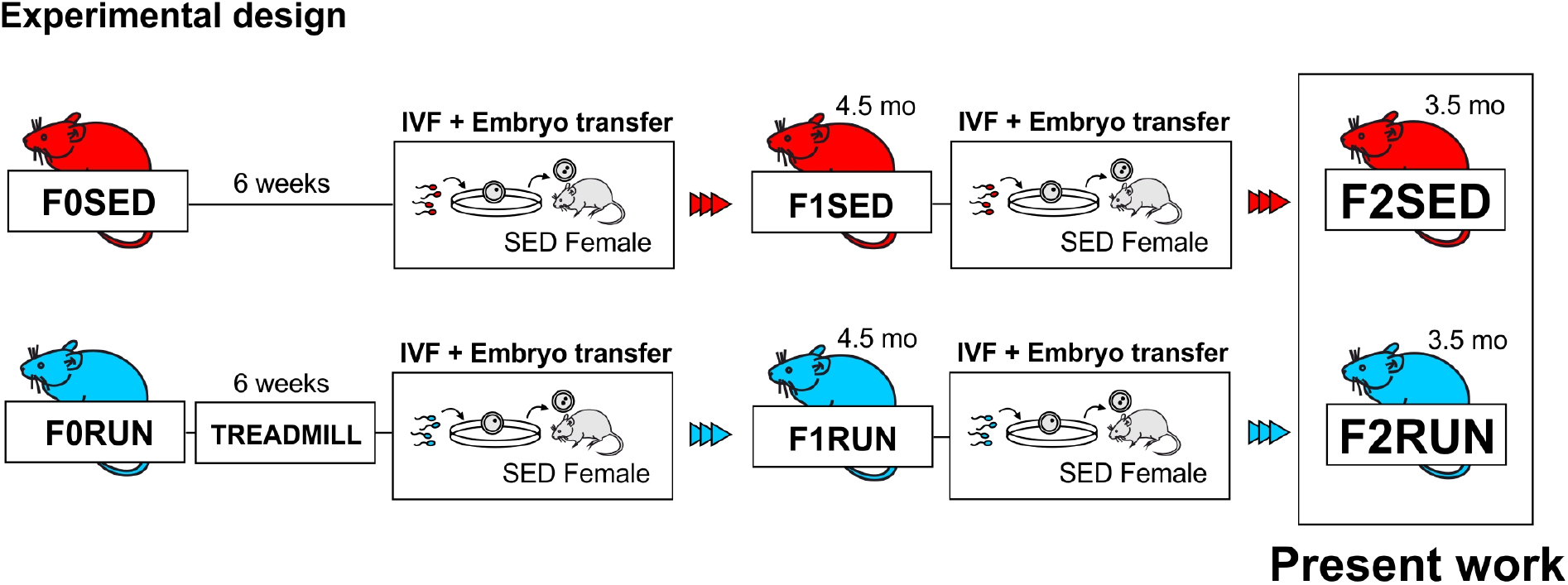
Experimental design. Mice were derived from sedentary (F0SED) or exercised (F0RUN) grandfathers. F0RUN animals were submitted to treadmill exercise of moderate intensity during 6 consecutive weeks, while F0SED animals remained in home-cage without any access to wheel or treadmill. At the end of these 6 weeks, their sperm was collected and used to produce offspring (F1SED and F1RUN animals, respectively) via *in vitro* fertilization (IVF) methods. Both F1 groups lived until 4.5 months of age, when their sperm was collected. F1 animals remained sedentary through the entire period. Then, a new offspring was generated via IVF (F2SED and F2RUN animals). These litters were analyzed from 3.5 months of age, when behavioral tests were conducted. At the end of the behavioral testing, animals were sacrificed for histological and epigenetic analysis. Both groups (F2SED and F2RUN) animals also remained sedentary through the entire experiment.

**Figure 2.**
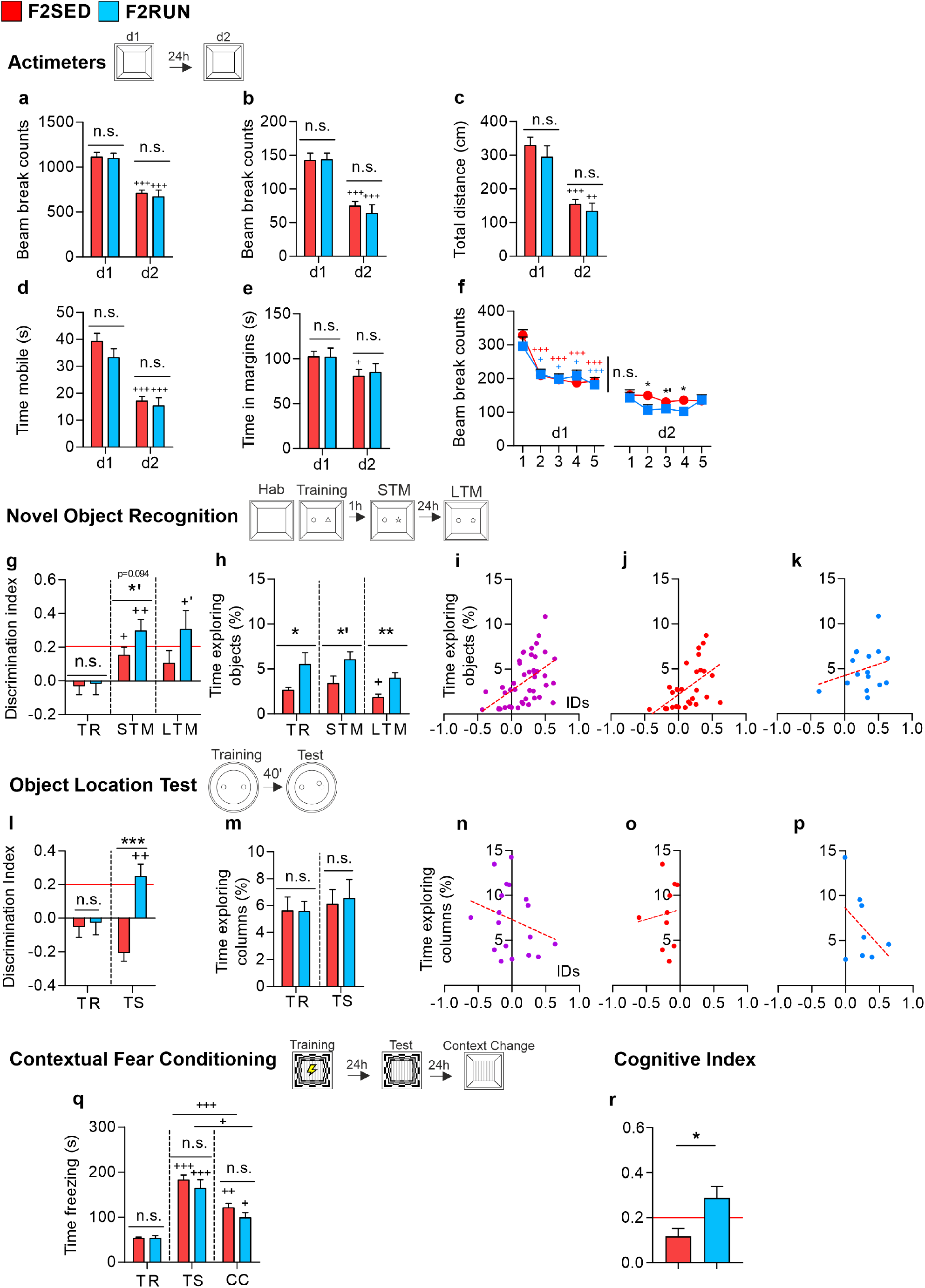
Behavioral analysis. **a-f,** Basal activity. F2 animals (F2SED, n=15; F2RUN, n=8) were subjected to activity measurements in activity cages for 5 min. A scheme of the actimeter test is shown. Both groups showed similar levels of horizontal activity (**a**) in a novel environment (day 1, d1) and a known environment (day 2, d2). Also, both groups showed a significant decrease in activity in the known environment compared to the novel environment. These were also true for the vertical activity (**b**), total distance traveled (**c**) and time in movement (**d**). Both groups spent similar time in the margins of the cage (**e**), with no significant differences between them in d1 or d2. F2SED animals spent significantly less time in the margins on the second day of the test. Horizontal activity was also checked minute by minute (**f**). Both groups showed a normal pattern of activity, with high exploration time during the first minute and a gradual, significant decrease of exploration the resting four minutes. On the second day of the test (d2), F2RUN animals showed significant less exploration in some minutes. **g-k**, Novel Object Recognition test (NOR). A scheme of the test is shown. Evaluation of discrimination indexes (DIs, F2SED, n=15; F2RUN, n=8) reports a better performance of F2RUN animals (**g**). Groups did not have a preference for any object in the training phase (TR), as they had DIs proximal to zero. In the short-term phase (STM), F2RUN animals showed a trend for a better discrimination than F2SED animals, but only the mean DI of F2RUN animals fulfilled the standard criteria to consider that discrimination has occurred (DI>0.2, red line). In the long-term phase (LTM), F2RUN animals also showed slightly higher DIs compared to F2SED animals (not significant). F2RUN also showed a trend to a higher DI compared to the training phase, while F2SED did not. Also, in the LTM phase only F2RUN animals fulfilled the standard criteria. Time exploring objects was also checked (**h**). F2RUN animals showed higher levels of exploration in all three phases. Correlational analysis between exploration time and DIs in STM and LTM phases was performed either in all F2 animals (**i**, showing a positive and significant correlation -Spearman’s rho, ρ=0.597, p<0.001-, or by group, **j,k**). Only F2SED animals showed a significant positive correlation between time exploring objects and DIs (**j**, Spearman’s rho, ρ=0.654, p<0.001), no correlation was found in F2RUN animals (**k**, Pearson’s coefficient, r=0.276, p=0.3). These results indicate that the higher exploratory behavior of F2RUN animals do not explain their better performance in NOR. An Object Location test (OL) was applied to check pattern separation abilities (**l-p**). A scheme of the test is shown. Evaluation of DIs (F2SED, n=10; F2RUN, n=8) reports a better performance of F2RUN animals (**l**). Both groups did not prefer any of the columns in the TR phase. In the test phase (TS), F2RUN animals had significantly higher DIs than the F2SED group, and fulfilled the standard criteria for discrimination. In this test, groups showed similar levels of exploration in both phases (**m**). The correlational analysis did not show a positive relation between time exploring columns and DIs in this test (**n**). This was also true for analysis by group (**o,p**). A Contextual Fear Conditioning (CFC) was applied to study memory and discrimination abilities in an emotional-aversive context (**q**). A scheme of the test is shown. Both groups (F2SED, n=13, F2RUN, n=8) had similar levels of freezing in the Training phase (TR), showing similar abilities for contextual classic conditioning. They also exhibited similar levels of freezing when were placed again in the conditioning chamber 24h later (TS). For both groups, the time freezing in TS was significantly higher compared to TR, as expected. In the Context Change phase (CC), animals were placed again in the conditioning chamber but with different walls (plain white vs checkers pattern). In this phase, both groups showed context discrimination, as indicated by a significant reduction in freezing levels compared to TS phase, although the time freezing in CC was significantly higher compared to TR in both groups, as expected. A cognitive index (**r**) was calculated to summarize cognitive performance. We can see how only F2RUNs show a CI above 0.2, having a significantly higher cognitive performance than F2SED. All data are shown as mean ± SEM. For comparisons between independent groups, *=p<0.05, **=p<0.01, ***=p<0.001; *’= 0.05>p<0.099. For comparisons between dependent groups, +=p<0.05, ++=p<0.01, +++= p < 0.001; +’= 0.05>p<0.09.

Next, we analyzed cognition using the novel object recognition test (NOR), the object location test (OL), and the contextual fear conditioning (CFC). In NOR we used a version specifically designed to make it difficult to distinguish the replacement of a known object by a novel one as previously described (Fontán-Lozano et al., 2007) (McGreevy et al., 2019). We found that while the control F2SED did not meet the discrimination criteria (Figure 2g, see definition below), the F2RUN mice fulfilled the criteria for short-term (1 h, STM) and a trend for long-term (24 h, LTM) memory tests (DI higher than 0.2, and training *vs* test comparisons significantly different in STM (mixed ANOVA F_(1,21)_=8.035 p=0.01, n_p_^2^=0.277; *post-hoc* comparisons adjusted by Bonferroni p=0.01) and a trend in LTM (p=0.066)). F2RUN animals explored both the known and novel objects longer than the F2SED (Figure 2h, Mann-Whitney U test, training U=21, p=0.025; STM U=27, p=0.076; LTM U=15, p=0.006). To test if the differences in exploration time could explain the differences observed in the DIs, a correlational analysis between exploration time and DIs in STM and LTM phases was performed in all F2 animals. Indeed, it may be relevant because a longer exploration time correlates with higher DIs when considering all animals (Figure 2i, Spearman’s rho, ρ=0.597 p<0.001), but this correlation is only significant for F2SED animals DIs (Figure 2j, Spearman’s rho ρ=0.654 p<0.001) and not for F2RUN mice (Figure 2k). These results indicate that the higher exploratory behavior of F2RUN animals cannot explain their better performance in NOR.

In the object location test, only F2RUN animals significantly discriminated the small change in the spatial location of the objects in a difficult version of the test (Figure 2l, related sample t-test, t(7)=-4.198 p=0.004). In this case, no significant differences in exploration time were found (Figures 2m-p).

In the contextual fear-conditioning test, an emotional-aversive test, we found no significant differences between experimental groups in neither the acquisition of the task nor the change in behavior after distinguishing a different context (Figure 2q). Thus, both groups under study learned and recalled the task to the same extent.

To check that the results were comparable and replicable between OL and NOR, we elaborated a global cognitive index, considering the test phase of OL, and both the STM and the LTM phase (NOR). Only the F2RUN animals significantly discriminated both the change in spatial location of objects and the objects themselves on the difficult version of both tests (Figure 2r, t-test, t(21)=-2.828 p=0.005).

Next, we characterized the adult hippocampal neurogenesis of the animals by a battery of histological markers in the dentate gyrus. We found no significant differences in neural stem cell numbers as measured by Sox2/GFAP labelling (Figure 3a,b), nor in cell proliferation in the granule cell layer (Figure 3c,d) as judged by pH3 labelling, nor in cell progenitor number / differentiating, immature newborn neurons measured by DCX/CLR (Figure 3e,f). As expected, no significant changes were found in the area occupied by the dentate subgranular layer (Figure 3g).

**Figure 3.**
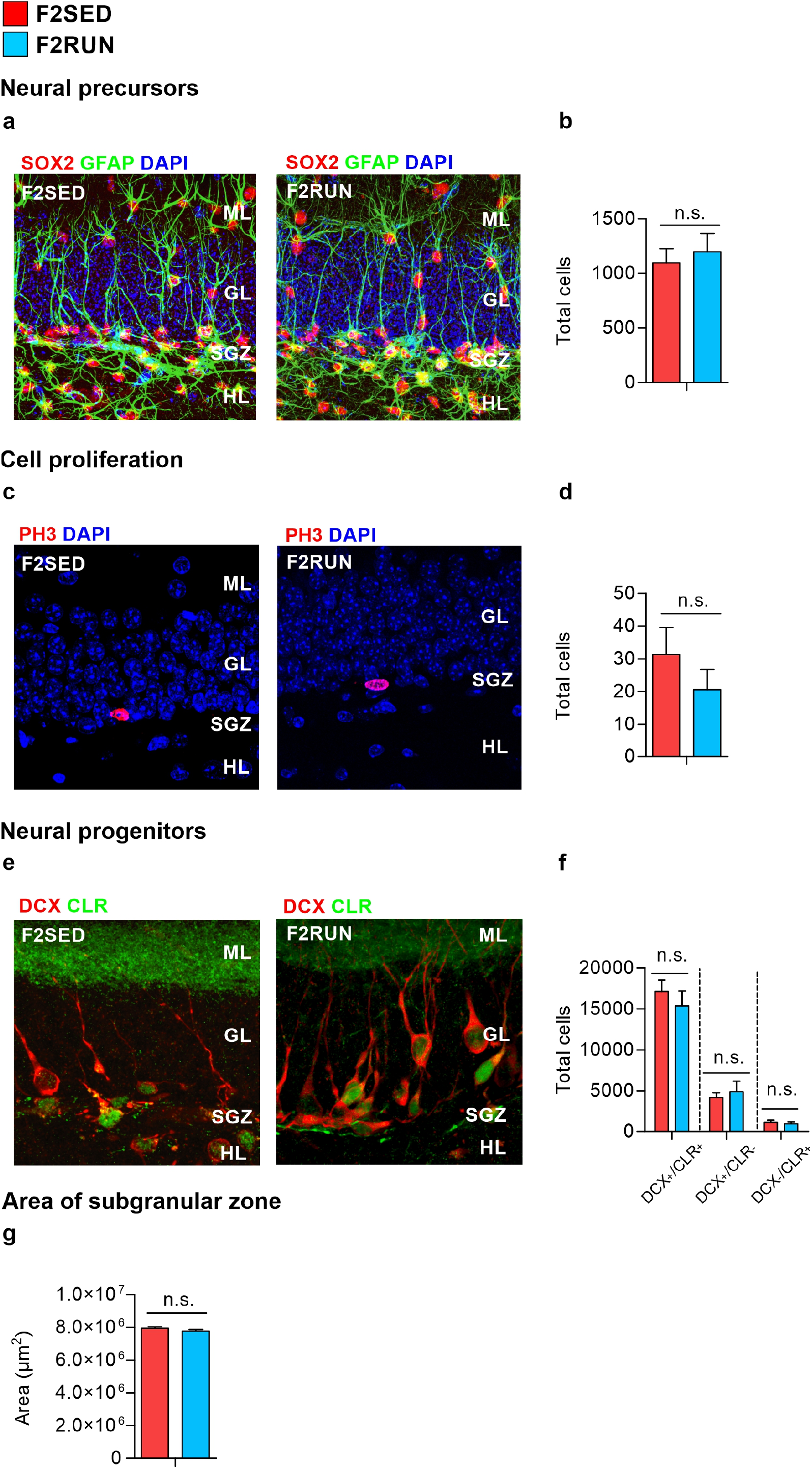
Histological analysis. Immunohistochemistry against a battery of markers was used to characterize the adult hippocampal neurogenesis (AHN). Total cell numbers were estimated by stereological unbiased protocols (F2SED, n=13, F2RUN, n=8). Mean total cell numbers are shown. Neural stem cells/precursors (**a,b**) were analyzed by the Sox2^+^/GFAP^+^ cell number in the subgranular zone of the hippocampal dentate gyrus (**a**). No significant differences between experimental groups were found (**b**). Cell proliferation (**c,d**) in the dentate gyrus granule cell layer was assessed by pH3 immunohistochemistry. pH3^+^ cells shown in **c** were counted, but no significant differences between groups were found (**d**). The immature, differentiating cell subpopulation in the granule cell layer (**e,f**) showed no significant differences in DCX^+^/CLR^-^, DCX^+^/CLR^+^, and DCX^-^/CLR^+^ cell subpopulations (**f**, respectively from left to right). Similarly, no differences were found in the subgranular zone area (**g**).

Finally, we analyzed the differential expression of microRNAs by a smallRNAseq of the F2 hippocampus (n=6 per group). We found 35 significantly differentially expressed (sDE) microRNAs, 15 microRNAs with a lower expression and 20 microRNAs with a higher expression in exercised F2 compared to sedentary F2 (Figure 4a). Relative homogeneity of the differential microRNA expression of both experimental groups can be observed when the heatmap is clustered according to the cognitive index (Figure 4a). The global expression change can be observed in a volcano plot (Figure 4b) with indication of fold change and statistical significance to appreciate which are the microRNAs with their relative differential expression, as well as the most represented gene ontology categories (Figure 4c). Interestingly, we have found that the expression level of two microRNAs significantly, negatively correlates with the F2 cognitive index: 298_5p (R −0.64, p=0.025), and 144-v1_5p/144-v2_5p (R −0.68, p=0.016), the lower the expression levels of these miRNAs induced by the grandparents exercise, the better the cognitive index of F2 litters (Figure 4d,e). The overlapping target genes of these two miRNAs appear in GO annotations “GOCC_SYNAPSE”, “GOCC_NEURON_PROJECTION”, and “GOBP_NEUROGENESIS”. Finally, we compared the results of the F2 smallRNAseq with our previous results of RNAseq of both F0 and F1 (after GSEA analysis of the RNAseq and the Molecular Signatures Database collection MIR: microRNA targets, or DAVID analysis), to find that the target gene set of 11 of the 35 sDE microRNAs in F2 had been also enriched in F0 and F1, while the target genes of another 3 of the 35 were found also enriched specifically in F1 (GSEA analysis, Figure 5). Some of the sDE microRNAs (Figure 4) presents both 3p and 5p arms coexpressed and apparently functional (for that reason they add up to 35). Instead, in figure 5 each microRNA is represented only one time, thereby adding 27. Moreover, the target genes of 6 of those 35 F2 sDE microRNAs had been differentially expressed (sDEGs, DAVID analysis) also either in F0 or F1 (table 1, here all the co-expressed, co-regulated forms are represented, thereby adding 9 different forms).

**Figure 4.**
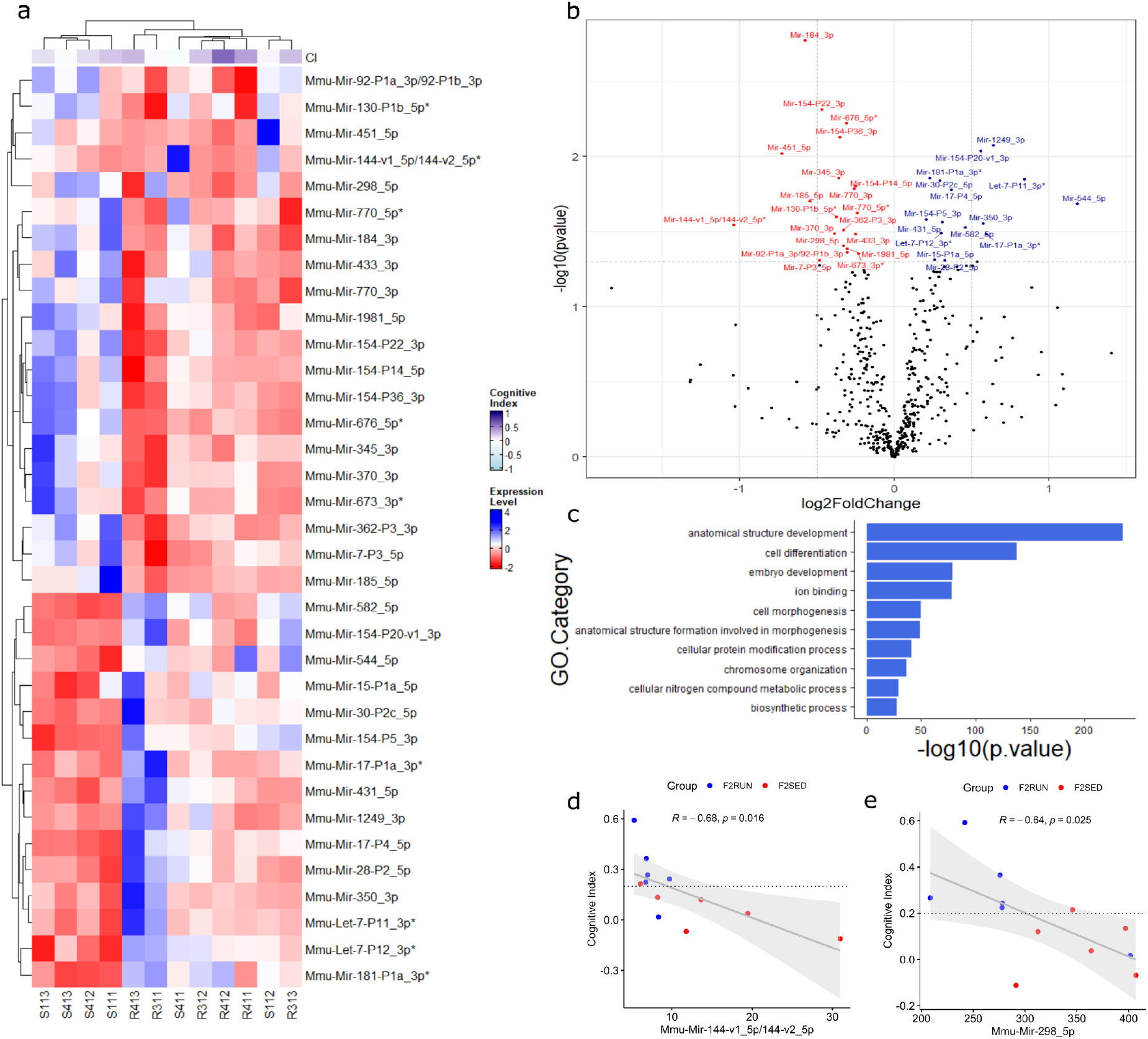
smallRNA analysis. miRNA-seq differential expression analysis between the F2 mice, grandsons of either sedentary or exercised grandfathers. **a,** Clustered heatmap of differentially expressed microRNAs between the 2 groups (p <0.05). Samples labelled with an “R” are from grandsons of exercised grandfathers and samples labelled with an “S” are from grandsons of sedentary grandfathers. All samples have been grouped by unsupervised hierarchical clustering (Pearson’s correlation) based on the Cognitive Index (CI) as a cognitive performance measure. **b,** Volcano plot showing the global expression change across the 2 groups. microRNAs with a p-value less than 0.05 and positive log2 fold change (upregulated) are represented as blue dots. microRNAs with a p-value less than 0.05 and negative log2 fold change (downregulated), as red dots. The x-axis represents the log2-transformed fold change of microRNA expression (Runner/Sedentary) and the y-axis is the p value (-log10-transformed). **c,** Gene Ontology (GO) Over Representation analysis (ORA) on the differentially expressed miRNAs gene targets. Bar plot shows the top 10 enriched GO terms with a p<0.05. **d-e**, significant negative correlation (Pearson’s) plots between cognitive index and microRNA expression levels of microRNAs 298_5p (R −0.64, p=0.025), and 144-v1_5p/144-v2_5p* (R −0.68, p<0.01).

**Figure 5.**
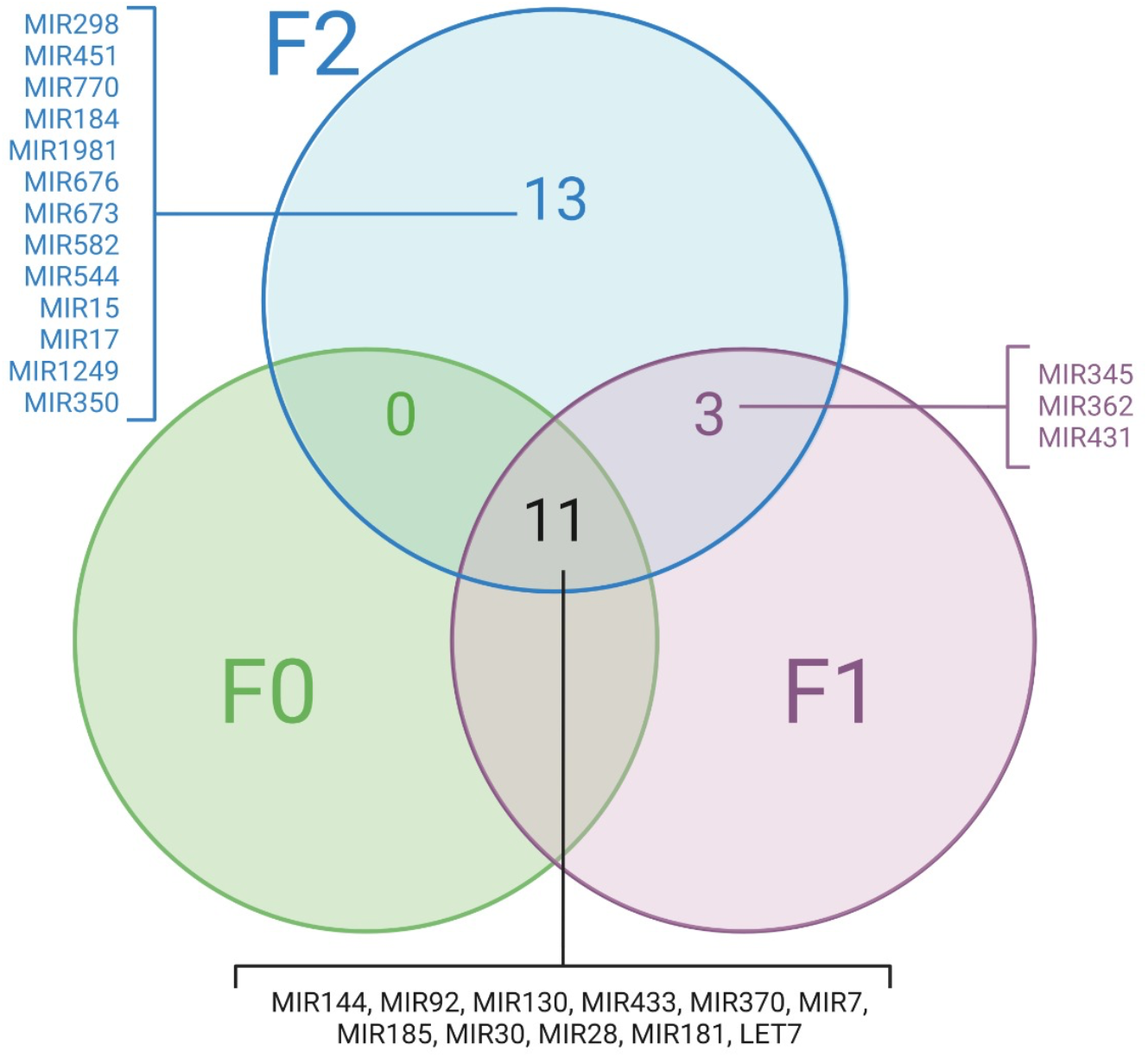
Comparison of F2 smallRNAseq results with F0-F1 RNAseq to reveal whether the putative target genes of those microRNAs significantly, differentially expressed in F2 were already differentially expressed in F0 and/or F1. The gene sets from “Molecular Signatures Database collection MIR: microRNA targets” enriched after a GSEA analysis of F0 and F1 RNAseq were used to detect coincidences with the F2 sDE miRNAs. We found the number of genes represented in the Venn diagram for each of the generations. 11 F2 miRNAs or its target genes in F0 and F1 appeared consistently different across 3 generations. This diagram shows 27 microRNAs because each sDE miRNA from the figure 4 is represented here just one time for clarity, while the figure 4 shows 35 because all the different sDE miRNA arms are represented (for example, figure 4 represents both 3p and 5p arms of some miRNA when both forms have been found sDE, while figure 5 represents one time that miRNA). Venn diagram licensed by BioRender.

**Table 1.** List of F2 sDE miRNAs whose target genes are sDEGs in the RNAseq of F0 or F1 (DAVID analysis). 6 microRNAs were found. 3 of them appeared with both arms co-regulated, thereby 9 different forms are represented in this table.

| **<u>miRNA</u>** | **<u>Gene Target</u>** | **<u>Differentially expressed in</u>** | **<u>Gene description</u>** |
| --- | --- | --- | --- |
| 144_3p | Srsf10 | F1 | Serine/arginine-rich splicing factor 10 |
| 144_3p | Hnrnpl | F1 | Heterogeneous nuclear ribonucleoprotein L-like |
| 1981_5p | Vipr2 | F0 | Vasoactive intestinal peptide receptor 2 |
| 1981_5p | Pnrc2 | F1 | Proline-rich nuclear receptor coactivator 2 |
| 370_3p | Hnrnpl | F1 | Heterogeneous nuclear ribonucleoprotein L-like |
| 544_5p | Pou4f1 | F0 | POU domain, class 4, transcription factor 1 |
| 582_5p | Irx2 | F0 | Iroquois homeobox 2 |
| 770_3p | Zmiz2 | F1 | Zinc finger, MIZ-type containing 2 |
| 770_5p | Chrna6 | F0 | Cholinergic receptor, nicotinic, alpha polypeptide 6 |

## DISCUSSION

The behavioral phenotype and AHN markers expression were analyzed in the second generation from either sedentary or exercised male mice (patrilineal design). Only the generation F0 (grandfathers) experienced moderate, forced physical exercise training, while the F1 and the F2 (the latter, analyzed here) were all sedentary. Grand-offspring (F2SED) from sedentary F0 animals were compared to those (F2RUN) from exercised F0 mice.

F2RUN mice learnt to distinguish a novel object from a known object 1 h after training, and were able to recall this information after 24 h, in a difficult non-spatial NOR design which F2SED animals were not able to accomplish fully. Independently, F2RUN animals spent longer times exploring both objects. However, this specific (object driven) exploratory activity does not significantly correlate with F2RUN DIs (while it is the case for F2SED). On the other hand, the fact that F2RUN animals significantly explored the open field for slightly shorter times on day 2 (Figure 2f) in the actimeters may suggest memory of an already known arena. Together, these results show that neither the different unspecific, spontaneous exploratory activity nor a specific (object-driven) higher exploratory activity in F2RUN animals can account for the higher discrimination abilities between known and novel objects. This points to a specific improved ability in F2RUN mice to distinguish a difficult object replacement, apart from an increased interest in exploring the environment. These findings are similar to those reported in intergenerational experiments analyzing the first generation (McGreevy et al., 2019).

F2RUN animals discriminated better than F2SED mice in a spatial, difficult version of the object location test, by fulfilling the discrimination criteria while the controls did not. Also, control mice did match the criterion for the immobile object (the one in its previous location), thereby displaying no preferences by either object, suggesting these animals did not appreciate any change in the environment. The results in both tasks (NOR and OL) points to a general improvement in both spatial and non-spatial abilities caused by the intervention. Again, similar results have been reported previously in the first generation (McGreevy et al., 2019).

It is relevant to consider that the experiments conducted here were designed to be sensitive to the difficulty level, although the findings might also be task-dependent. Certainly, both difficulty level- and task-dependent results have been found in intergenerational experiments with interventions like environmental enrichment (Benito et al., 2018) and physical exercise (McGreevy et al., 2019). Here we have shown, first, no significant differences between groups in a different cognitive task like contextual fear conditioning (test trial), and second, that a salient change in the features the animals use to distinguish the contexts in CFC (easy versions) reveals no differences (change of context trial). Nevertheless, the aversive stimulus in the CFC, a repeated electric shock, is quite reinforcing. Although it has been previously shown the intergenerational inheritance of exercise-induced effects on conditioned fear (Short et al., 2017; Yeshurun & Hannan, 2019), we have found no differences in F2 generation (transgenerational).

We found no differences in the adult neurogenesis rate in the hippocampal dentate gyrus, neither in neural stem cell number, nor in proliferation, nor in immature newborn neuron numbers. This result is not striking because this niche has largely reported as highly sensitive to lifestyle interventions (Llorens-Martín, 2018), and a paradigmatic example of metaplasticity (García-Segura, 2009)(Llorens-Martín et al., 2009). It is not surprising that a highly sensitive process like AHN responds to physical activity (for a recent review see for example (Bettio et al., 2020)). This response is intergenerationally inherited by the next generation (F1, (Benito et al., 2018)(McGreevy et al., 2019)) but is no longer present transgenerationally, in a generation (F2) whose fathers (F1) were sedentary as well (present results). Nevertheless, it cannot be ruled out that more subtle changes in the neurogenic subpopulation of the hippocampus (like synaptic boutons, for example) might be altered by the intervention but not detected by the analyses performed here or in previous works, as suggested by the GO annotation (see below) of the sDE miRNAs found in the present work.

Previous studies have found that the procognitive effects of physical exercise rely on changes on different miRNAs (Benito et al., 2018; Goldberg et al., 2021 PMID 34625536). Therefore, to explore potential mediating mechanisms of these results, we performed a smallRNAseq of hippocampus of adult F2SED and F2RUN animals. We also compared the results of this smallRNAseq with the RNAseq of F0 and F1 (McGreevy et al., 2019). We have found 35 significantly differentially expressed (sDE) miRNAs (either over- or under-expressed between F2SED and F2RUN), involved in relevant functions like structure development, cell differentiation, and ion binding, among others. This finding is especially interesting after comparing to the sDEGs in F0 and F1. As the target genes of many miRNAs may be either directly regulated by a specific miRNA or indirectly by a complicated network of different, coregulatory group of miRNAs, we made a comparative analysis of the target genes of F2 sDE miRNAs compared to the F0 and F1 GSEA enriched “MicroRNA targets”. Eleven of the 35 F2 sDE miRNAs are associated to target gene sets found also enriched in both F0 and F1 (GSEA analysis), pointing to a small group of active miRNAs involved across generations in the determination of the traits analyzed here. Moreover, specific target genes of 6 sDE miRNAs in F2 were also sDEGs either in F0 or in F1 after DAVID analysis, pointing to a group of specific genes involved in this transgenerational inheritance. Especially noticeable is the miRNA 144, one of the two sDE miRNAs significantly correlating to cognitive index in F2 (the other is 298). miRNA 144 is found in both F0 and F1 list of enriched microRNA target genes (McGreevy et al., 2019), and two of its target genes are also sDEGs in F1 (Srs10 and Hnrnpl). Finally, GSEA analysis revealed that target genes of miRNAs 144 and 298 are involved in the biological function Neurogenesis and the cellular components Synapse and Neuron Projection. Together, these findings suggest common pathways of epigenetic/genetic regulation leading to the phenotype described here.

Our findings are relevant because, to our knowledge, only adverse outcomes have been largely reported to be transgenerationally inherited in mammals (Gapp et al., 2014)(Bohacek & Mansuy, 2015)(Jawaid et al., 2018)(Cunningham et al., 2021)(Pang et al., 2021). On the contrary, only intergenerational inheritance of beneficial, pro-cognitive effects have been described after physical-cognitive activity so far, (Benito et al., 2018)(McGreevy et al., 2019)). Here we describe a transgenerational inheritance of the positive effects of moderate, forced physical exercise training in male mice on cognition. We found that some but not all the effects intergenerationally inherited by the first generation of sedentary litters (McGreevy et al., 2019) are transgenerationally inherited in the second generation (present results), pointing to a partial vanishing of the influence of the parental exercise when the lifestyle intervention is removed across generations. The grandfather exercise-induced cognitive improvement of grandsons is revealed in both spatial and non-spatial tests and seems to be task-specific and difficulty-level sensitive. A small group of miRNAs and target genes is strongly suggested as a potential epigenetic/genetic mechanism accounting for this effect. These findings might also be interpreted as suggesting that sedentary lifestyle effects (adverse body and brain health effects) can be transmitted to next generations.

Together, these results point to an unexpected heritability of the beneficial effects of a moderate exercise program on cognition, and clear the way to explore further the molecular mechanisms mediating these effects (like for example specific microRNAs reported here) that might be used as pharmacomimetics of healthy lifestyles. Nevertheless, these findings may be valuable data supporting evidence-based health policies in contexts like development, adult disease, and aging.

## MATERIAL AND METHODS

### Animals

All animals from the three generations (F0, F1 and F2) were housed under standard laboratory conditions, in accordance with European Union Directive 2010/63/EU. All experiments were performed according to the European Community Guidelines (Directive 2010/63/EU) and Spanish Guidelines (Royal Decree 53/2013) and related norms, and they were first validated by the Committee of Ethics and Animal Experimentation of the Cajal Institute (20/05/2016), subsequently favorably evaluated by the CSIC Ethics Committee (Subcommittee of Ethics) of the Spanish Research Council (07/27/2016) and eventually authorized by the competent authority, the Animal Protection Area of the Department of Environment of the Community of Madrid (10/26/2016 and 06/19/2020).

### F0 male progenitors (grandfathers of F2 animals)

Animals were purchased and housed individually as in McGreevy et al., 2019. They were randomly assigned to the experimental conditions (sedentary vs exercised). The exercised group (F0RUN) underwent the exercise protocol for 6 weeks while the sedentary group (F0SED) remained in the home cage. The exercise protocol was implemented as described elsewhere (McGreevy et al., 2019). After treatment, sperm of these animals was used to obtain F1 males via *in vitro* fertilization.

To analyze whether transgenerational exercise-driven effects were germline dependent, the progeny of both F1 and F2 was generated through in vitro fertilization (IVF) and embryo transfer by using the male sperm of F0 and F1 respectively. After birth, the litter sizes were homogenized between (foster) mothers, and after weaning, the subjects from different experimental groups were randomly mixed (Bohacek & Mansuy, 2015).

### F1 male progenitors (fathers of F2 animals)

F1SED animals were obtained from sperm of F0SED progenitors and F1RUN animals were obtained from sperm of F0RUN progenitors using *in vitro* fertilization. On postpartum day 21 (P21), a mixed weaning strategy was used to avoid group effects and to minimize possible litter effects due to cohabitation. At 4.5 months old, their sperm was used to obtain F2 animals via IVF. Both F1SED and F1RUN males remained sedentary during all the procedures.

### F2 males (current generation)

F2SED (n=15) and F2RUN males (n=8) were obtained from two F1SED or F1RUN progenitors respectively, using *in vitro* fertilization methods. On postpartum day 21 (P21), a mixed weaning strategy was used to avoid group effects and to minimize possible litter effects due to cohabitation. Behavioral testing started at 3.5 months of age. Animals were sacrificed at 4.5 months old.

### *In vitro* Fertilization

IVF and embryo transfer were conducted at the Mouse Embryo Cryopreservation Facility of the National Centre for Biotechnology, Spanish National Research Council as described elsewhere (McGreevy et al., 2019).

### Behavioral Assessment

#### Activity assessment

A VersaMax Legacy Open Field activity box (Omnitech Electronics) was used to study locomotor activity. F2 animals underwent a two-day protocol (5 min in the activity cage per day). On the first day of the protocol, the spontaneous locomotor activity was measured for 5 minutes in a novel arena. The following day, animals were placed in the same arena for another 5 minutes. Behavioral measures included total horizontal activity, horizontal activity per minute, total vertical activity, total distance moved, total time mobile and total time in margins and center.

#### Novel Object Recognition (NOR)

A NOR protocol was applied in an open wooden box of 42 x 32 x 31 cm. The test was applied as described in previous work (McGreevy et al., 2019). It consisted of four phases: habituation (Hab), training (TR), short-term memory evaluation (STM) and long-term memory evaluation (LTM) (Figure 2). In the habituation phase, the animals were left to freely explore the box without any object (5 min). During the training phase, each animal was placed in the arena containing two different objects in the center of the box (objects A and B) and was allowed to explore them freely for 5 minutes. Next, each animal was removed and put back in the home cage. One hour after training, animals were placed back in the NOR arena containing a familiar object (object A) and a novel object (object C) and allowed to explore them for 10 minutes. At the end of this phase, all animals were placed in their home-cage. Twenty-four hours after training, animals were tested again for 10 minutes in the arena containing the familiar object (A) and a novel object (object D). The box was cleaned with 0,03% acetic acid solution between trials. Time exploring each object was manually scored. Discrimination indexes were calculated using the next formula for TR (in STM and LTM the time exploring B is replaced by either C or D, respectively):

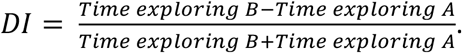

#### Object Location Test (OL)

OL test protocol was applied in a PVC circular arena (35 centimeters in diameter and 20 centimeters high) as described in (McGreevy et al., 2019). The test consisted in two phases (Figure 2): training and test. In the training phase, animals were left to explore the arena for 4 minutes. Two equal plastic columns were placed symmetrically along the diameter, at the same distance from the walls. At the end of the phase, animals were placed again in their home-cage. Forty minutes after the beginning of the training phase, animals were put back in the circular arena for 4 minutes (test phase). One of the columns was displaced. The arena was cleaned with 0,03% acetic acid solution between trials. Time exploring each column was manually scored. Discrimination indexes were calculated using this formula:

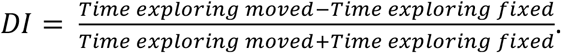

In OL, the column displacement distance determines the difficulty of the test. The shorter the displacement distance the more difficult the test. In this work we used only difficult versions of this test.

#### Cognitive Index (CI)

This index was calculated to summarize the cognitive ability and to compare overall performance in both spatial and non-spatial tests, both short and longterm trials, between experimental groups. Cognitive index was calculated as the mean of the DI scores of the three tests:

#### Contextual Fear Conditioning (CFC)

A fear conditioning apparatus (Ugo Basile Fear Conditioning 2.1, 46003 Mouse Cage) was used to test contextual aversive memory and context discrimination abilities. The test consisted of three phases: Training (TR), Test (TS) and Context Change (CC). In the training phase (5 minutes), animals were left to freely explore the conditioning chamber (17×17×25cm) with walls with checkers pattern (context A) for 3 minutes. At minute 3:00, 3:30 and 4:00, a floor-shock (0.5mA, 2 seconds of duration) was administered through the floor grid. The animal was left another minute in the cage before the ending of the test. Test phase was conducted 24h after the training phase. Animals were put back in the conditioning chamber with checkers pattern (context A) and left to explore for 5 minutes, no shock applied. 24h after the test phase, the CC phase was performed. The animals were placed again in the conditioning chamber but with white walls (context B). Freezing time was automatically scored with ANY-MAZE (V6.0).

#### Tissue collection

Animals were deeply anaesthetized with 10 mg/kg bw of pentobarbital (Euta-Lender) and transcardially perfused with 0.9% saline. Next, the brain was removed and dissected down the midline into 2 hemispheres. The left hemisphere was fixed by immersion in 4% paraformaldehyde in phosphate buffer (PB) for 24 hours at room temperature. The following day, each hemisphere was washed with PB and stored at 4°C.

#### Histology

Serial coronal brain sections (50 μm of thickness) containing hippocampal formation were obtained from each hemisphere on a Leica VT1000S vibratome and individually collected in a 96-multiwell plate filled with PB 0.1M. Plates were kept at 4°C until further analysis.

#### Immunohistochemistry

Each series of sections used for immunohistochemical analysis were composed by systematic sampling of the hippocampus in the rostro-caudal axis (8-9 hippocampal sections per animal, 50 μm of thickness each section, 400 μm apart from each other). For single or double staining, slices were initially preincubated in PB with 1% Triton X100 and 1% bovine serum albumin (BSA) (PBT-BSA). Primary antibodies (DCX, Goat anti-doublecortin, 1:500, Santa Cruz; SOX2, Goat anti-sex determining region Y-box 2, 1:200, R&D Systems; CLR, Rabbit anti-calretinin, 1:3000, Swant; GFAP, Rabbit anti-glial fibrillary acidic protein, 1:2000, Abcam; pH3, Rabbit anti-phospho-histone H3, 1:500, Millipore) were incubated in agitation with PBT-BSA for 1h at room temperature and 72h at 4°C. Secondary antibodies (Donkey anti-goat alexa fluor 594, Donkey anti-rabbit alexa fluor 594, Donkey antirabbit alexa fluor 488, Donkey anti-rat alexa fluor 594; Invitrogen, 1:1000) were incubated with PBT-BSA for 1h at room temperature and 24h at 4°C. Cell nuclei were counter-stained with 4’,6-diamino-2-phenylindole (DAPI, Sigma-Aldrich, 1:1000).

#### Stereology

The number of neural precursor cells was estimated using 7 physical disectors per animal, obtained with confocal microscopy (Leica TCS SP5, oil immersion 40× objective). Disectors were randomly positioned in rostro-caudal sections of the granule cell layer of the dentate gyrus visibly containing the SGZ. Neural precursor cells were identified following these criteria: 1) SOX2^+^/GFAP^+^ staining; 2) Cell body positioned in SGZ; 3) Radial glia-like morphology with a long process across the GCL; and counted using Image J (Fiji 1.46). Density per disector and the total number of neural precursors along hippocampus was calculated in the same way than below described for DCX/CLR cells.

Phosphohistone H3^+^ (Ph3^+^) cells were identified and counted for each animal in a complete series of hippocampal sections using a fluorescence microscope (Leica DMI 6000 B). The total number of positive cells was estimated by the optical fractionator method, multiplying the total cell number by the sampling fraction (i.e., 1/8 of every hippocampal section of an animal was counted, thus the sampling fraction was 8). Only cells stained for pH3^+^, positioned in subgranular zone and showing mitotic morphology were counted.

Doublecortin and/or calretinin expressing cells (DCX^+^/CLR^+^, DCX^+^/CLR^-^ and DCX^-^/CLR^+^ cells) were counted using Image J (Fiji 1.46) on 7 physical disectors obtained with confocal microscopy (Leica TCS SP5, oil immersion 63× objective) in each animal. Disectors were randomly positioned in rostro-caudal sections of the granule cell layer of the dentate gyrus visibly containing de subgranular zone (SGZ). Density of DCX^+^/CLR^+^, DCX^+^/CLR^-^ and DCX^-^/CLR^+^ cells was calculated per each disector. The total number of each type of cell per animal was estimated multiplying the cell density by the area of the SGZ of each animal. The area of the SGZ was estimated for each animal using the Cavalieri method. The length of the SGZ was measured in a different series of Nissl staining with Neurolucida software. This length was then multiplied by the distance between 2 systematic-sampled slices (400 μm).

#### miRNA analysis

Hippocampus regions were dissected and immediately frozen at −80°C. miRNA isolation was performed using microRNA micro kit (Qiagen, Qiagen BgmH, Germany) using manufacturer protocol. Briefly, samples were homogenized in Qiazol Lysis reagent, cleaned by sepharose column and eluted in RNase free water. Total RNA quality was assessed using vertical electrophoresis (QSep100, Bioptic Inc) and cleanness and concentration was evaluated using Nanodrop One (Thermofisher). Samples were remitted to Novogene for microRNA isolation and sequencing. Libraries were prepared using NEB Next^®^ Multiplex Small RNA Library Prep Set for Illumina^®^ (Set 1) (Cat No. E7300). Libraries were then sequenced on the Illumina^®^ Novaseq6000 to generate 50 bp single reads according to the manufacturer’s protocol. The volume of data was 10 Million reads equivalent to 0.5 Gb of data. The quality score of at least 85% of bases was > Q30 (Phred Quality Score 30).

After quality checking with FastQC (Wingett & Andrews, 2018) and miRTrace (Kang et al., 2018) software, the reads were analysed using miRge3.0 v0.1.2 (Patil & Halushka, 2021), aligning against the MirGeneDB database. Normalization of miRNAs raw counts and differential expression analysis were performed with DESeq2 R package (Love et al., 2014). The software DIANA-miRPath v.3.0 (Vlachos et al., 2015) was used to associate differentially expressed miRNAs to target genes (with Tarbase, v7.0) in order to determine associated and potentially affected GO terms and KEGG metabolic pathways.

For the miRNA transgenerational coincidences, we first searched for every one of the 35 sDE F2 miRNAs on the list of miRNAs whose target genes were sDEG in F0 or F1 (11 were both sDE on F2 and their target genes were sDEGs on F0 and F1, 3 were sDE on F2 and their target genes were sDEGs only on F1). Then we consulted every target gene of all 35 sDE miRNAs (via mirdb.org (Chen & Wang, 2020)) and filtering by a threshold of 80 on the target score) on F2 and run the list against every sDEGs gene on the RNA-seq. 9 genes were found either on F0 or F1. Lastly, we submitted every gene target of mir-144 and mir-298 to GSEA (Subramanian et al., 2005) to compute overlaps, checking collections M1, M2, M3, M5, M8 and MH. We found the following enriched gene sets: MIR_298_5P, MIR_144_5P, MIR_451A, MIR_6951_3P, GOCC_SYNAPSE, GOCC_NEURON_PROJECTION, MIR_466D_5P, MIR_466K, GOBP_NEUROGENESIS and MIR_466I_5P.

#### Statistical analysis

For comparisons between two independent groups, the T-test was applied when the dependent variable was normally distributed. The Mann-Whitney U Test was used in the case of non-normal distribution. For comparisons between two dependent groups, paired sample t-test was applied if the dependent variable was normally distributed. When not, Wilcoxon Signed Ranked Test was used. For behavioral tests that contained between-subject and within-subject measures with three levels, Mixed ANOVA was applied when met the normality and homogeneity assumptions. If the normality and/or the homogeneity of variances assumptions were violated, a Friedman test was used followed by a post hoc Wilcoxon Signed-Ranked Test. To study correlations, Pearson correlation was calculated when data followed normal distribution. Spearman correlation was used with non-normal distributions. All data were analyzed using SPSS Statistics (IBM, v.26.0.0). Data are shown as mean ± SEM. To test normality, the Shapiro-Wilk Test was applied. Extreme values identified by the software were eliminated from the analysis. Inter-group differences: *p<0.05, **p<0.01, ***p<0.001. Trends in inter-group differences 0.05 ≥ *’p < 0.099. Within-group differences: +p<0.05, ++p<0.01, +++p<0.001. Trends in intra-group differences 0.05 ≥ +’p < 0.099. All graphs were created in GraphPad Prism 5.

## Notes

### Competing Interest Statement

The authors have declared no competing interest.

